# Discovery and fine-mapping of kidney function loci in first genome-wide association study in Africans

**DOI:** 10.1101/2020.06.09.142463

**Authors:** Segun Fatumo, Tinashe Chikowore, Robert Kalyesubula, Rebecca N Nsubuga, Gershim Asiki, Oyekanmi Nashiru, Janet Seeley, Amelia C Crampin, Dorothea Nitsch, Liam Smeeth, Pontiano Kaleebu, Moffat Nyirenda, Stephen Burgess, Nora Franceschini, Andrew P Morris, Laurie Tomlinson, Robert Newton

## Abstract

Genome-wide association studies (GWAS) for kidney function have uncovered hundreds of risk loci, primarily in populations of European ancestry. We conducted the first GWAS of estimated glomerular filtration rate (eGFR) in Africa in 3288 Ugandans and replicated the findings in 8224 African Americans. We identified two loci associated with eGFR at genome-wide significance (p<5×10^−8^). The most significantly associated variant (rs2433603, p=2.4×10^−9^) in *GATM* was distinct from previously reported signals. A second association signal mapping near *HBB* (rs141845179, p=3.0×10^−8^) was not significant after conditioning on a previously reported SNP (rs334) for eGFR. However, fine-mapping analyses highlighted rs141845179 to be the most likely causal variant at the HBB locus (posterior probability of 0.61). A trans-ethnic GRS of eGFR constructed from previously reported lead SNPs was not predictive into the Ugandan population, indicating that additional large-scale efforts in Africa are necessary to gain further insight into the genetic architecture of kidney disease.

## Introduction

Chronic Kidney Disease (CKD) is a global public health problem, with adverse outcomes of kidney failure, cardiovascular disease, and premature death. Unfortunately, little is known about the genetic epidemiology of CKD in continental Africa where the disease is at least three times more frequent [1]. Given the central importance of Africa to our human origins, there is a clear scientific and public health need to develop large-scale efforts examining genetic diversity and disease susceptibility across diverse populations within Africa [2]. The marked genomic diversity and allelic differentiation among populations in Africa, in combination with the substantially lower linkage disequilibrium (LD) among genetic variants, provides excellent opportunities to gain new insights into disease aetiology and genetic fine-mapping that have relevance for all ancestry groups. However, despite the value of conducting such research in Africa, there is no known genome-wide association study (GWAS) of kidney function in continental Africa, with published studies of African ancestry individuals being limited to African Americans [3,4]. Whilst African Americans typically have a large proportion of African ancestry, several studies have shown that the genetic architecture of African Americans is distinct from that of Africans from continental Africa [5]. The African American population reflects admixture of people of West and central-west African descent, adding to the relevance of studying populations from other regions of Africa.

Here, we conducted the first continental African GWAS of estimated glomerular filtration rate (eGFR), a measure of kidney function used to define CKD, including 3288 individuals from Uganda in East Africa. Subsequently, associations were validated through GWAS of eGFR in 8224 African Americans from the Women’s Health Initiative (WHI). Together, these GWAS comprised a total of 11,512 African ancestry individuals, which we used to: (i) identify loci associated with eGFR; (ii) fine-map loci by taking advantage of the finer-grained LD pattern of African ancestry populations; and (iii) evaluate the predictive power of an eGFR genetic risk score (GRS) into the Ugandan population that was derived from previously-reported trans-ethnic lead SNPs.

## Results

### Discovery genetic association analysis

The characteristics, quality control and imputation of the 3288 Ugandan participants from the General Population Cohort (GPC) are presented in the Methods. We analysed associations of eGFR for 20,594,556 SNPs that met a minor allele frequency (MAF) threshold of at least 0.5% in a merged panel of imputed GWAS and whole-genome sequences. We tested for association in a linear mixed model implemented in GEMMA, which accounted well for population structure and relatedness as shown by the genomic inflation factor (λ) ~1.01. We identified two loci attaining genome-wide significance (*p*<5×10^−8^) in GPC (Table 1, Figure 1): *GATM*(lead SNP rs2433603, p=1.0 x 10^−8^) and *HBB* (lead SNP rs141845179, p = 3.0 x 10^−8^). Both loci have been previously reported as associated with eGFR [6]. *GATM* encodes a mitochondrial enzyme that belongs to the amidinotransferase family. This enzyme is involved in creatine biosynthesis, whereby it catalyzes the transfer of a guanido group from L-arginine to glycine, resulting in guanidinoacetic acid, the immediate precursor of creatine. The haemoglobin beta (*HBB*) gene provides instructions for making a protein called betaglobin. Beta globin causes sickle cell anemia. Absence of beta chain causes beta-zero-thalassemia, and reduced amounts of detectable beta globin cause beta-plus-thalassemia.

**Table 1:** Description of Meta-analysis genome-wide significant loci

| Table 1: Description of Meta-analysis genome-wide significant loci |  |  |  |  |  |  |  |  |  |  |  |  |  |
| --- | --- | --- | --- | --- | --- | --- | --- | --- | --- | --- | --- | --- | --- |
|  |  |  |  |  |  | Uganda |  |  |  | WHI |  | Meta-Analysis |  |
| Locus | Lead SNP | Chr | BP (b37) | EA | NEA | Beta | SE | EAF | Pvalue | Pvalue | EAF | Pvalue | N |
| *GATM* | rs2433603 | 15 | 45646226 | T | C | 0.145 | 0.025 | 48% | $1.0 \times 10^{-08}$ | $7.3 \times 10^{-04}$ | 53% | $2.4 \times 10^{-09}$ | 11512 |
| *HBB* | rs141845179 | 11 | 5244665 | G | T | 0.266 | 0.048 | 8% | $3.0 \times 10^{-08}$ | NA | N/A | NA | 3288 |

**Figure 1a:**
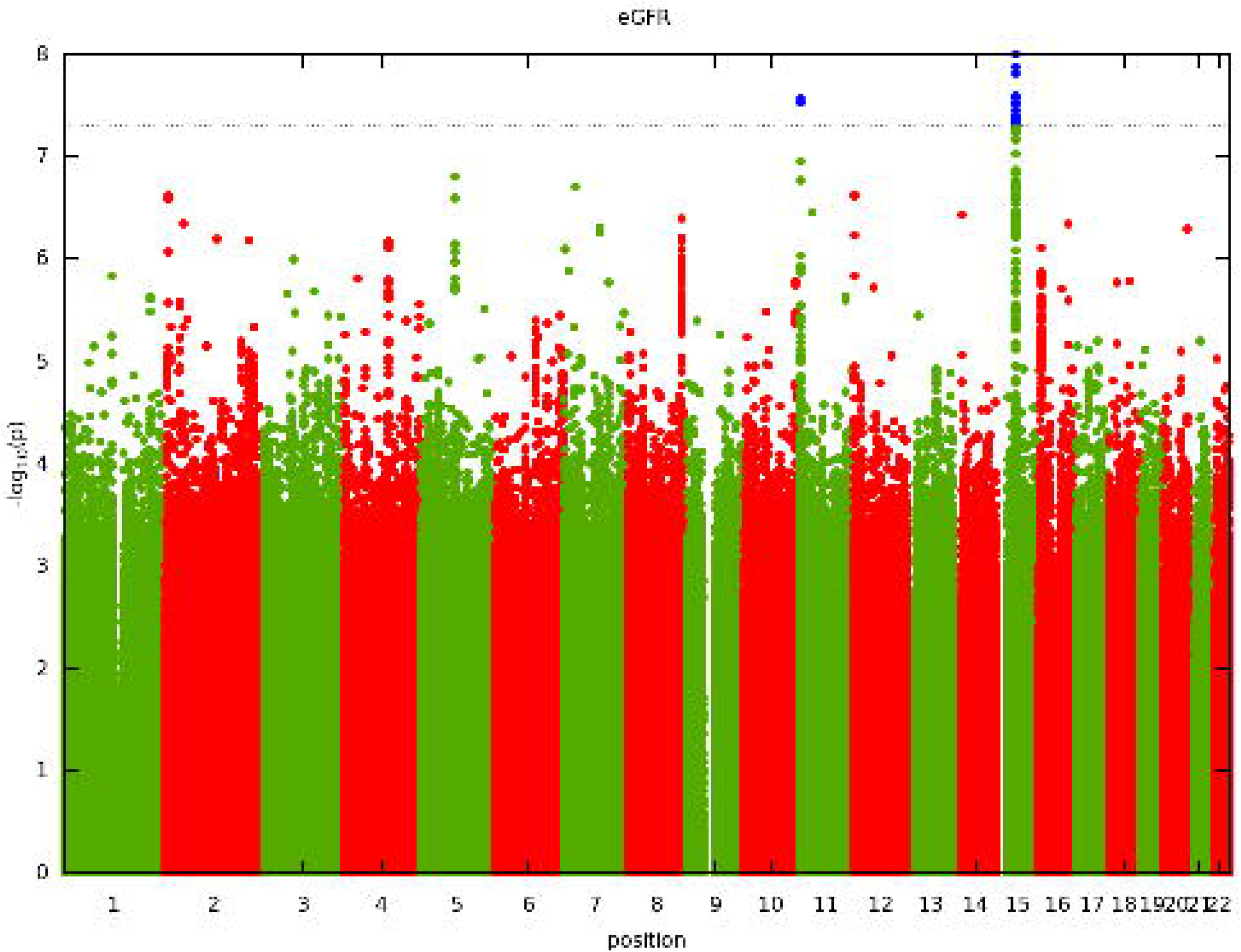
Manhattan plot of genome-wide associations of eGFR. Each point denotes a variant, with the X-axis representing the genomic position and Y-axis representing the association level −log 10 (P-value) for the association of all variants with MAF >=0.005. The dotted line shows the genome-wide significant P-value of 5×10^−8^.

**Figure 1b:**
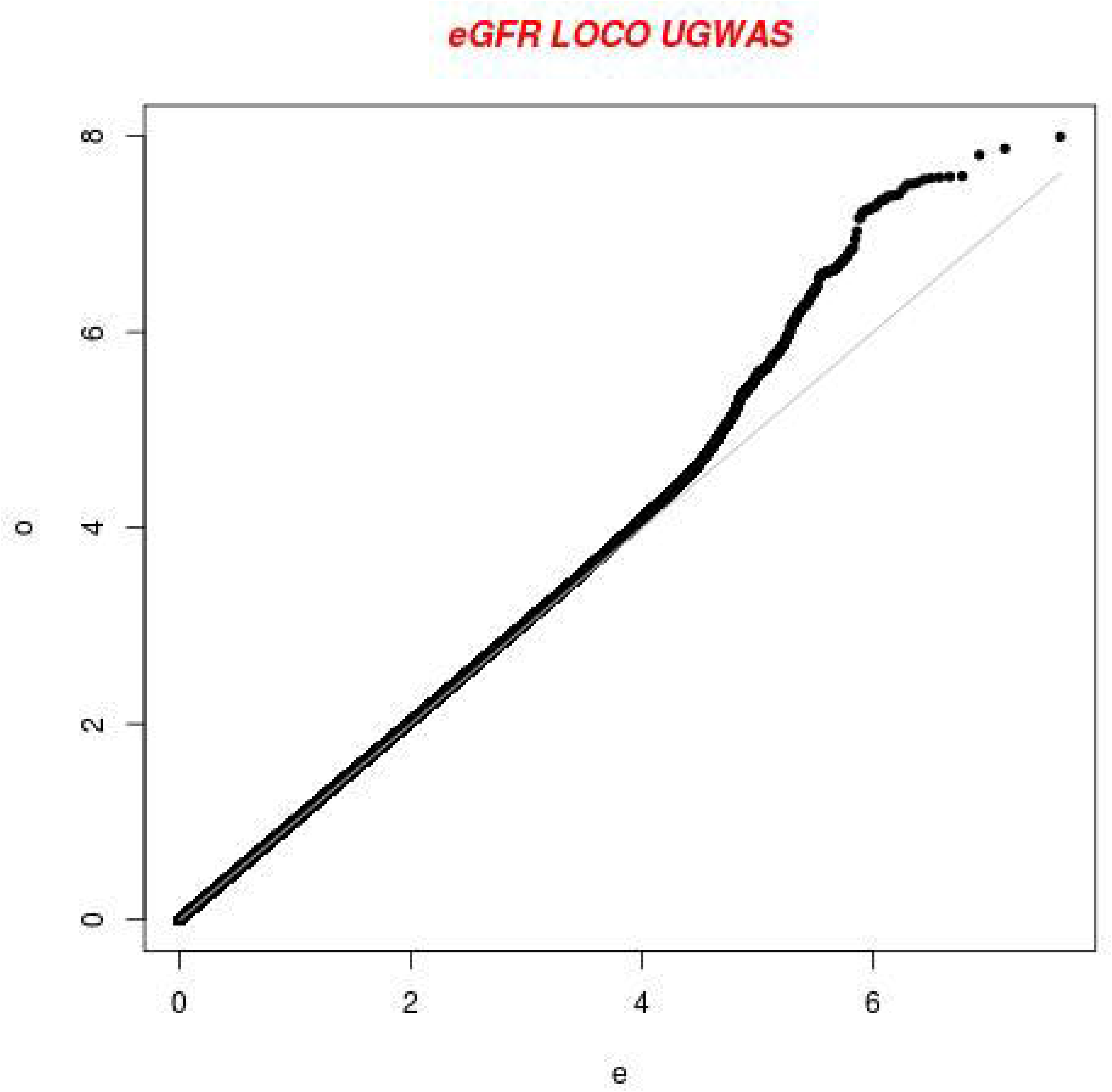
Visually inflated QQ plot of genome-wide associations of eGFR with an inflation factor of 1.01.

**Figure 2:**
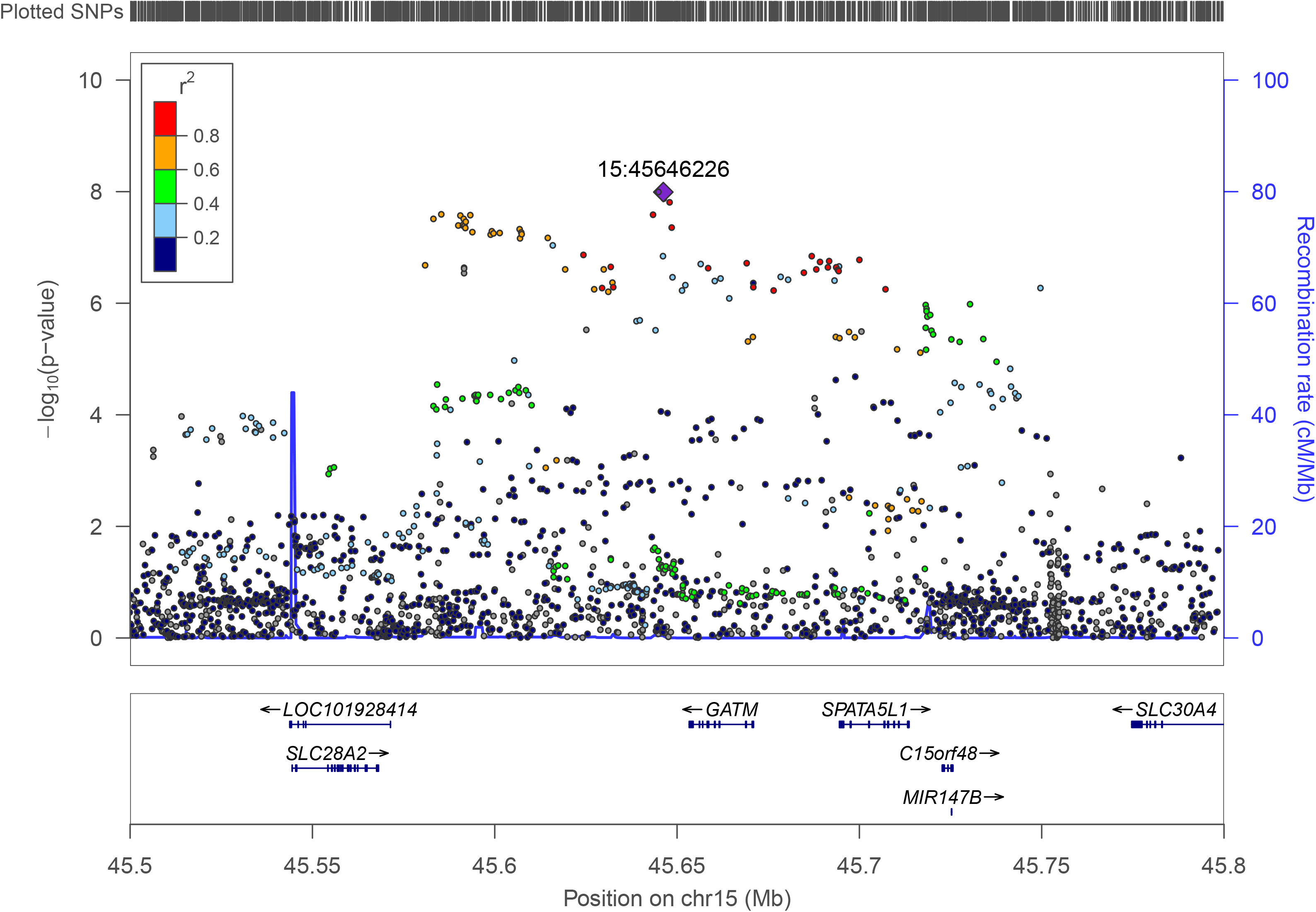
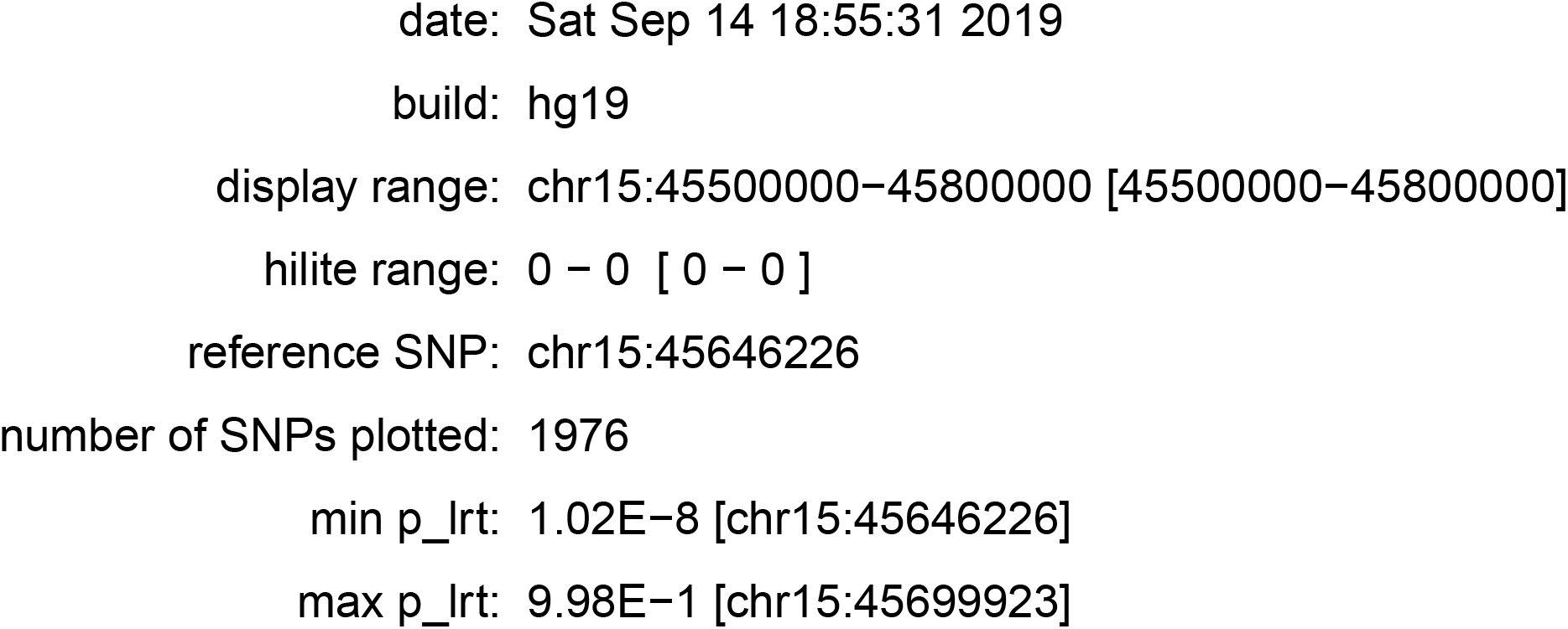
Regional association plot for eGFR in 3288 non-missing Uganda individuals. The lead SNP rs2433603 (15:45646226) (p = 1.01 × 10-8) located in in *GATM* on chromosome 15 is labelled and coloured in purple. LD (r2) was calculated based on the Ugandan SNP genotypes used in this study.5×10-8.

**Figure 3:**
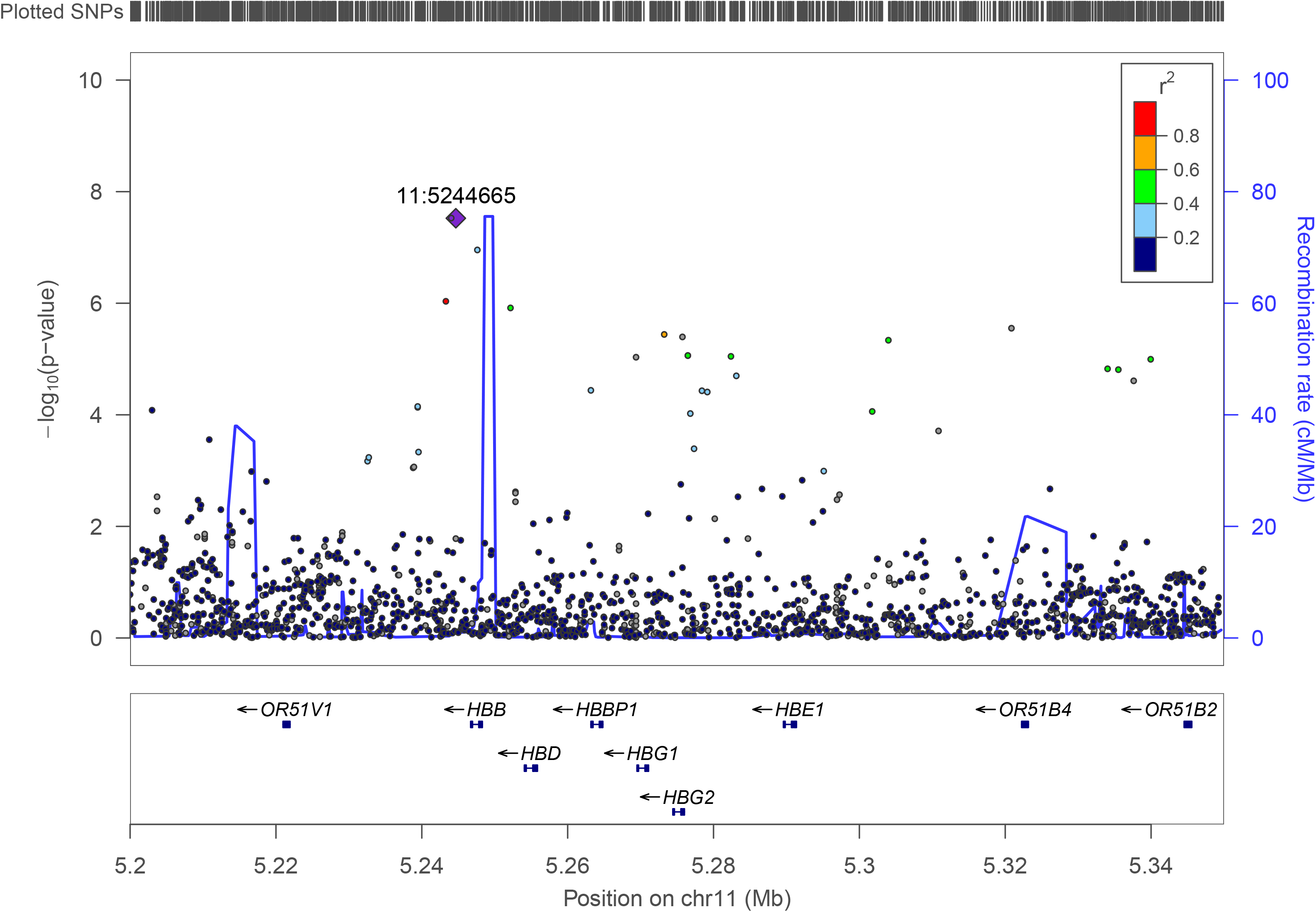
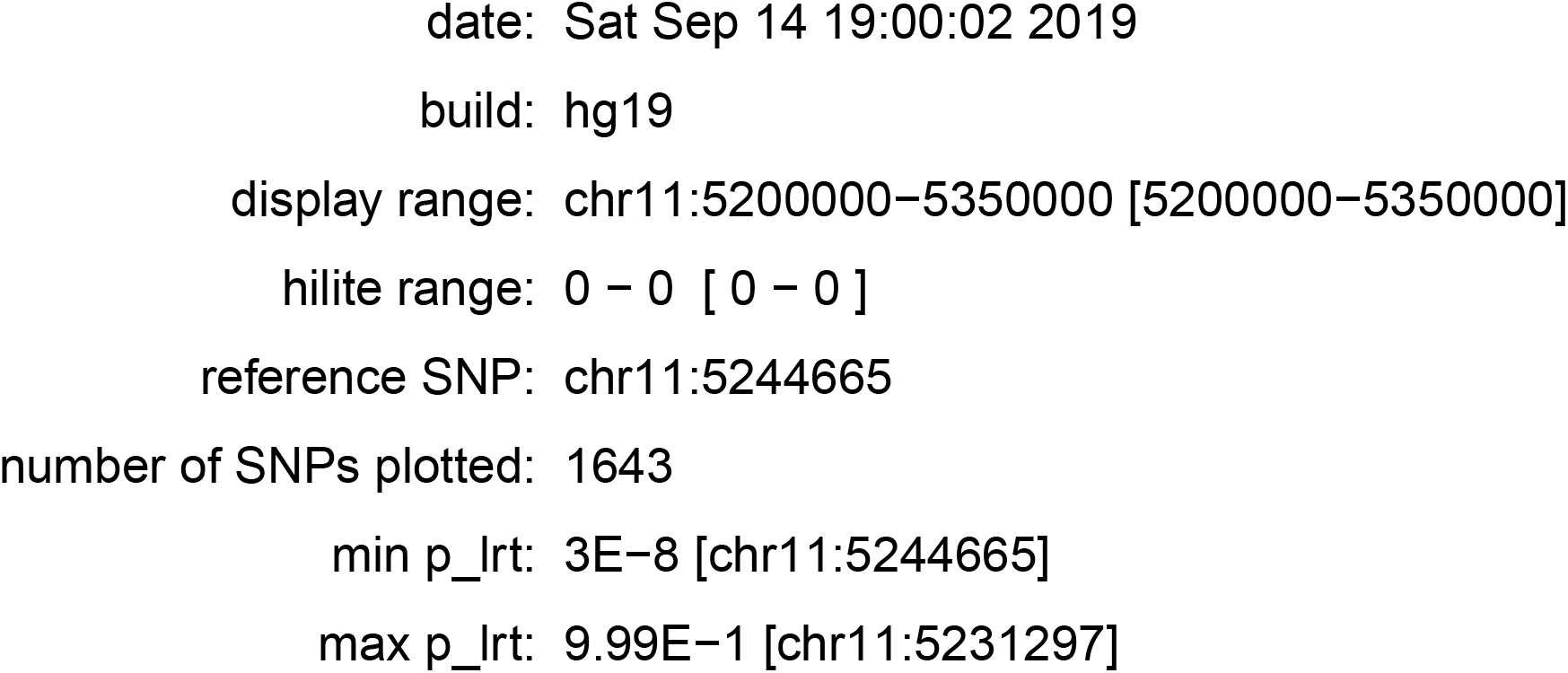
Regional association plot for eGFR in 3288 non-missing Uganda individuals. The lead SNP rs141845179 (11:5244665) (p = 2.99 × 10-8) located in *HBB* on chromosome 11 is labelled and coloured in purple. LD (r2) was calculated based on the Ugandan SNP genotypes used in this study.5×10-8.

To investigate the relationships between the signals in Uganda and those reported in other populations, we performed conditional analyses. The most significant signal for eGFR association at *GATM*, rs2433603, is distinct from previously reported associations (rs1145077, rs1153855, rs1145093) at this locus (conditional p = 4.0 x10^−7^, Supplementary figure 1). This genetic variant is monomorphic in European ancestry populations and rare in East Asian ancestry populations in the 1000 Genomes Project Phase 3 [19]. At the *HBB* locus, after conditioning on the previously reported SNP (rs334) at this locus, the association with the lead SNP (rs141845179) was no longer significant. The variant (rs334) in *HBB* has also been previously associated to other kidney traits, including urinary albumin to creatinine ratio (UACR) [24].

### Replication and Meta-Analysis

All 1309 SNPs showing strong evidence of association (p<5 x 10^−5^) in Uganda were considered for replication in WHI. In view of the different scales of the effect sizes in GPC and WHI (because of different measurements of eGFR), association summary statistics were aggregated across the two studies using the fixed-effects meta-analysis based on the sample size weighting of Z-scores (Table 1) in METAL [17]. We replicated the association signal at *GATM* in WHI (p= 7.3 x 10^−4^, meta-analysis p= 2.4 x 10^−9^). However, the lead SNP at the *HBB* locus was not available in WHI.

### Fine-mapping of loci attaining genome-wide significance

Fine-mapping of loci was undertaken in the region mapping 500kb up- and down-stream of the lead SNP. Fine-mapping of the *GATM* locus was performed using the meta-analysis of Uganda and WHI, whilst fine-mapping of the *HBB* locus was based only on Uganda since the lead SNP was not reported in WHI. For each locus, we constructed the 99% credible set of SNPs that together account for 99% of the posterior probability of driving the association. At the *GATM* locus, the 99% credible set consisted of 63 variants, and no variant accounted for more than 50% of the posterior probability (the lead SNP, rs1145092 had posterior probability of 13%). At the *HBB* locus, the 99% credible set consisted of 23 variants, with rs141845179 accounting for 61% of the posterior probability.

### Transferability of trans-ethnic eGFR GRS into Uganda

We used lead SNPs from a trans-ethnic meta-analysis [3] of eGFR to evaluate the predictive power of an unweighted GRS into unrelated individuals in the Uganda population (Table 2). We were unable to undertake a weighted GRS because different transformations of the trait were performed in the trans-ethnic meta-analysis and in the Ugandan GWAS. Whilst the GRS showed the correct direction of effect, it was not significantly associated with eGFR in the Uganda population (p=0.08) and accounted for only 0.04% of the trait variance in eGFR after accounting for age, sex and principal components to adjust for population structure.

**Table 2:** Regression coefficients for the association of the GRS and eGFR

| *Variable* | Model without GRS |  |  | Full Model |  |  |
| --- | --- | --- | --- | --- | --- | --- |
|  | Beta(SE) | P | R2 | Beta(SE) | P | R2 |
| *Age* | -0.92(0.01) | $< 2.0 \times 10^{-16}$ | | 0.93(0.015) | $2 \times 10^{-16}$ | |
| *Sex* | -2.53(0.56) | $7.9 \times 10^{-06}$ | | -2.55(0.53) | $6.8 \times 10^{-06}$ | |
| *PC1* | 22.62(21.53) | 0.29 |  | 21.77(21.53) | 0.31 |  |
| *PC2* | -40.46(22.01) | 0.07 | **0.5492** | -41.78(22.01) | 0.06 | **0.5496** |
| *PC3* | 32.57(20.17) | 0.11 |  | 33.18(20.16) | 0.09 |  |
| *PC4* | 24.34(23.62) | 0.30 |  | 24.62(23.61) | 0.29 |  |
| *PC5* | -2.95(21.05) | 0.89 |  | -3.40(21.04) | 0.87 |  |
| *GRS* |  |  |  | **-0.17(0.09)** | **0.08** |  |

#GPR

## Discussion

In the first GWAS of eGFR in continental Africa, we confirmed association with the previously-reported *HBB* locus. We also identified and successfully replicated a novel eGFR association that is distinct from previously reported signals at the *GATM* locus. Fine-mapping of the *HBB* locus identified a single variant, rs141845179, that accounted for 61% of the posterior probability of driving the association signal. This variant is non-coding, so would operate through regulation of gene expression, but is not a significant eQTL in kidney in GTEx or the Human Kidney eQTL Atlas [25]. This could be because the variant is rare in Europeans, and so not represented in these European-centric eQTL resources. This further strengthens the need for additional multi-omics resources in Africa.

The GRS derived from previously reported trans-ethnic lead SNPs for eGFR was not significantly predictive in the Ugandan population. This is mostly likely due to a lack of power in the Uganda GWAS because of small sample size. However, the lack of transferability could also be because of the way eGFR is calculated in continental Africa. Studies have shown that there is considerable potential error in the measurement of serum creatinine in continental Africa that might lead to inaccurate estimates of kidney disease at individual and population level [7, 26]. In this study, eGFR was calculated using the CKD-Epi formula, without use of the coefficient for African Americans [8]. The absence of a validated estimating equation for kidney function in Africans could be a contributing factor to the lack of GRS transferability. Another potential explanation for why the GRS was not significant is because the lead SNPs from the trans-ethnic analysis might not themselves be causal variants, and are not in LD with the causal variant in the Ugandan population because of differences in LD structure.

Despite the relatively high burden of CKD in Africa [9], there have been no previous GWAS of eGFR in continental Africa. One limitation of this first GWAS is that, with a small sample size, we are underpowered to reliably detect associations at genome-wide significance thresholds. We were also unable to apply more stringent threshold recommended for Africa population due to sample size [12]. However, our findings further highlight the importance of diverse ancestries for uncovering novel associations. Larger continental African metaanalyses are warranted to gain further insight on the genetic architecture of eGFR. In addition, while GWAS still remains a leading tool to identify loci contributing to complex diseases, to follow up significant findings and gain biological insights, the multi-omics resources that would inform these analyses need to be better represented in Africans. The study of populations in Africa provides a research framework to help characterise ethnic-specific patterns of variation in CKD among populations [10] and in a larger framework of studies, might also help identify population-specific genetic or environmental factors that may statistically interact with identified genetic loci. Given these scientific opportunities, the ascertainment and collation of genetic epidemiological resources with the statistical resolution to examine these associations in African populations is a high priority.

## Methods

### GPC Study participants

The recruited African individuals were part of the Uganda General Population Cohort (GPC), which is a population-based cohort of roughly 22,000 inhabitants around 25 neighbouring villages of Kyamulibwa, which is a subcounty of Kalungu district in the countryside in the south-west of Uganda. The cohort study was founded in the late 1980s by the Medical Research Council (MRC) UK in partnership with the Uganda Virus Research Institute (UVRI) to primarily investigate the trends in incidence and prevalence of HIV infection in Uganda. Samples were collected from research participants during a survey from the research study area. The study area is clustered into villages defined by governmental borders ranging in size from 300 to 1,500 dwellers and includes numerous families resident within households [11]. The GPC Round 22 study took place in 2011 through collaboration between the University of Cambridge, Wellcome Sanger Institute (WSI) and MRC/UVRI in Uganda. This study was approved by the Science and Ethics Committee of the UVRI, the Ugandan National Council for Science and Technology, and the East of England-Cambridge South NHS Research Ethics Committee United Kingdom. The study was contained within one annual survey round of the longitudinal cohort. The focus of the GPC Round 22 study was to investigate the genetics and epidemiology of communicable and non-communicable diseases to provide aetiological insights into the genetic variation in communicable and noncommunicable diseases.

### GPC Study design

The data collection of GPC Round 22 study consisted of five main stages, which took place in 2011 over the course of the year: mobilization (recruitment and consenting), mapping, census, survey, and feedback of results and clinical follow-up. The census consisted of a family questionnaire and questionnaire for the individual recruited from within the family. The family questionnaire was completed by the head of family or another responsible adult or emancipated minor member of the household. The household census questionnaire focused on sociodemographic information about the household, such as the quality of the house, property ownership, and employment of workers. The individual survey questionnaire captured information on members of a household including position within household, marital status, resident status, childbirth, and fertility, tribe, and religion. Information on lifestyle and health was obtained using a standard questionnaire. This included biophysical measurements and blood samples [11]. We genotyped 5,000 and sequenced 2,000 samples from 9 ethno-linguistic groups from the GPC.

### GPC Genotyping and quality control

5000 individuals were genotyped on the Illumina HumanOmni2.5-8 array. 4,872 were retained following a pre-quality control stage. GWAS genotype data were subjected to stringent quality control filtering. We considered a total of 2,314,174 autosomal variants, of which 39,368 were excluded because they did not pass SNP quality thresholds for call rate (< 97%, 25,037 SNPs) and deviation from Hardy-Weinberg equilibrium (HWE) (p < 10^−8^, 14,331 SNPs) as reported in [12]. We excluded a total of 91 individuals as they failed to meet the quality control cutoffs of call rate (>97%), heterozygosity in the range of mean±3SD, or because they failed the gender check criteria using the X-chromosome as a match. Three additional samples were also excluded because of high relatedness using identity by descent. We also excluded six genotyped samples that were found to be potentially contaminated.

### Curation of GPC Sequence Data

An additional 2000 Uganda samples (UG2G) underwent low coverage whole-genome sequencing on the Illumina HiSeq 2000 with 75bp paired end reads, at low coverage, with an average coverage of 4x for each sample. 1,978 of them passed QC. The workflow for data processing of UG2G has been previously described in more detail [12,15].

### GPC Haplotype phasing and imputation into genotype data

Haplotype phasing of GWAS data was carried out using SHAPEIT2 [20] with standard parameters. A previous study has shown that phasing with SHAPEIT2 in this cohort with dense genotype data provides very high accuracy even when pedigree structure is not explicitly specified during phasing [21].

Imputation of the pre-phased genotype data was carried out with IMPUTE2 [22] using a merged reference panel of the whole genome sequence data from the African Genome Variation Project [12], the UG2G described earlier and the 1000 Genomes Project phase 3 (1000 Genomes Project Consortium, 2015) following standard recommendations. Imputation was carried out in chunks of 2 MB and then concatenated. Imputed SNPs were further filtered for info quality >0.3 and a minor allele frequency (MAF) >0.5%. All duplicated variants were also removed.

### Merging of GPC genotype and sequence data

The final dataset used for this analysis included merged genotype data on 4,772 and sequence data on 1,978 individuals. We note that there are 343 individuals who have been genotyped and sequenced; for these individuals, we only included the sequence data, and not the genotype data. The final dataset, therefore, included 6,407 individuals (4,429 with genotype, and 1,978 with sequence data).

Following merging, we assessed and removed any systematic differences between imputed genotype data and sequence data. We did this by carrying out principal component analysis using merged data for the 343 individuals who had been genotyped and sequenced in duplicate to examine whether there was separation by data mode (imputed genotype data and sequenced data).. Full details were reported in [12]. For GWAS analyses, we only included a subset of variants (n = 20,594,556) that met a MAF threshold of at least 0.5%.

### GPC Laboratory Test and Phenotype definition

Creatinine was measured using the enzymatic method traceable to an isotope dilution mass spectrometry method (IDSM) [13]. Collectively, the serum creatinine level was measured in 3288 Uganda individuals for Round 22. The eGFR was calculated using the CKD-Epi formula, without use of the coefficient for African Americans [8]. We carried out the inverse rank normal transformation of eGFR residuals after adjusting for age, age^2^ and sex.

### Statistical methods for association analysis in GPC

GWAS was performed using the standard mixed model approach implemented in genomewide efficient mixed-model association (GEMMA) version 24 [14] for analysis of pooled data from 3288 individuals in GPC. In order to maximise discovery, we used the leave one chromosome out (LOCO) approach for analysis [15]. In this approach each chromosome is excluded from generation of the kinship matrix in turn, for association analysis for markers along that chromosome. This ensures that causal SNPs at a locus on a given chromosome are not used for generation of the kinship matrix used in analysis of that specific chromosome. Therefore, we generated 22 kinship matrices for analysis, each excluding the chromosome being analysed using the given matrix. For computational efficiency, and to avoid correlation effects due to LD, we LD pruned the data prior to calculation of the kinship matrix for each LOCO analysis.

For all loci attaining genome-wide significance that have been previously reported in GWAS of eGFR, we performed conditional analyses in GEMMA to determine whether the association signals were distinct. Specifically, we included genotypes under an additive model at previously reported lead SNPs as a fixed-effect. We also searched for evidence of multiple distinct signals of association in GPC by including genotypes at the lead SNP as a fixed-effect..

#### WHI Study Design

The Women’s Health Initiative (WHI) is a study of postmenopausal women and health outcomes. A total of 161 808 women aged 50–79 years old were recruited from 40 clinical centers in the United States between 1993 and 1998. Study protocols and consent forms were approved by the institutional review boards at all participating institutions. The WHI SHARe minority cohort includes 8515 self-identified African American women from WHI, who provided written informed consent for study participation and DNA analysis. The WHI African Americans tagged (WHI SHARe-A) is part of the WHI’s SNP Health Association Resource (SHARe), funded by the National Heart Lung and Blood Institute (NHLBI).

### WHI Genotyping, Imputation and Phenotype transformation

African American women who consented to genome-wide scanning (SHARe cohort-A) underwent genotyping with the Affymetrix Genome-Wide HumanSNP Array 6.0 containing 906,000 SNPs. The samples underwent initial quality control including removal of samples with poor DNA quality, abnormal sex chromosomes, relatedness, and low call rates. Additional quality control measurements were made at the SNP level assessing for Hardy-Weinberg Equilibrium (goodness-of-fit χ2> 10), call rates 90%, monomorphic SNPs, and minor allele frequencies 1%. We used frappe to estimate individual admixture, and estimates were included in models to account for population stratification. Full details of the quality control steps have been discussed in [3, 16].

As previously discussed in [3], after quality control, GWAS scaffolds were pre-phased and imputed using MaCH/minimac *r*^2^≥0.3 and 13,096,173 SNVs passing quality control were tested for association with eGFR. For each individual, eGFR was calculated from serum creatinine (mg/dL), with adjustment for age, sex and ethnicity, using the four variable the Modification of Diet in Renal Disease (MDRD) equation.

### Replication and Meta-analysis

All lead SNPs showing strong evidence of association (p<5×10-5) in Uganda was considered for replication in WHI. In view of the different scales of the effect sizes, association summary statistics of Uganda and WHI were aggregated using the fixed-effects meta-analysis based on the sample size weighting of Z-scores in METAL [17].

### Fine-mapping

We performed fine-mapping to identify potential causal variants for the locus ± 1MB of the lead SNPs in the *GATM* and *HBB* genes, using a Bayesian approach [23]. For *GATM*, this was meta-analysis but HBB was only Uganda. The Z score was then used to compute the Bayes factor for each SNP denoted as ***BF_i_***, given by

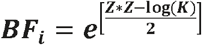

where K is the number of studies). The posterior probability of driving the association for each SNP was computed by

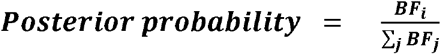

where the summation in the denominator is over all SNPs at the locus.

Ninety-nine percent credible set sizes were derived by sorting the ***BF_i_*** of the SNPs at the locus from the highest to the lowest, and then counting the number of SNPs needed to attain a cumulative posterior probability that is greater or equal to 0.99.

### Genetic Risk Score

First, we removed 817 first-degree relatives from the Uganda cohort derived from PIHAT values >0.5 and calculated principal components using –pca in PLINK [18]. Lead SNPs from a previously published trans-ethnic meta-analysis [3] of eGFR were selected and used to compute an unweighted GRS by counting the number of eGFR decreasing alleles, using the allelic scoring approach in PLINK [18].The predictive power of the GRS was evaluated by assessing the change in R2 when it was added to the linear model of the eGFR adjusted for age, sex and principal components.

## Acknowledgements

We thank all study participants and GPC staff who contributed to this study. S.F. received salary support from NIH grant U01MH115485 and the Makerere University-Uganda Virus Research Institute Centre of Excellence for Infection and Immunity Research and Training (MUII). MUII is supported through the DELTAS Africa Initiative (grant 107743). The DELTAS Africa Initiative is an independent funding scheme of the African Academy of Sciences (AAS), Alliance for Accelerating Excellence in Science in Africa (AESA), and supported by the New Partnership for Africa’s Development Planning and Coordinating Agency (NEPAD Agency) with funding from the Wellcome Trust (107743) and the U.K. government.TC is an international training fellow supported by the Wellcome Trust grant (214205/Z/18/Z).

This GPC is jointly funded by the UK Medical Research Council (MRC) and the UK Department for International Development (DFID) under the MRC/DFID Concordat agreement. Further funding was obtained from the Wellcome Trust (WT098051 and WT090132), the UK Medical Research Council and with federal funds from the National Cancer Institute, National Institutes of Health, under Contract No. HHSN261200800001E

The WHI program is funded by the National Heart, Lung, and Blood Institute, National Institutes of Health, U.S. Department of Health and Human Services through contracts HHSN268201600018C, HHSN268201600001C, HHSN268201600002C, HHSN268201600003C, and HHSN268201600004C.” The authors thank the WHI investigators and staff for their dedication, and the study participants for making the program possible. A full listing of WHI investigators can be found at: http://www.whi.org/researchers/Documents%20%20Write%20a%20Paper/WHI%20Investigator%20Long%20List.pdf”

NF is supported by the National Institutes of Health R01-DK117445, R01-MD012765, R21-HL140385. APM is supported by the National Institutes of Health R01-DK117445.

## Ethics Statement

This study was approved by the Science and Ethics Committee of the UVRI, the Ugandan National Council for Science and Technology, and the East of England-Cambridge South NHS Research Ethics Committee United Kingdom.

